# Upcycling banana peduncle fibers into mycelium-based composites for sustainable packaging and thermal insulation

**DOI:** 10.64898/2026.09.01.748491

**Authors:** Smruti B. Bhatt, Indumathi M. Nambi, Lakshminath Kundanati

## Abstract

The growing concerns due to plastic pollution in India have intensified the search for sustainable materials. Mycelium-based composites (MBCs) have emerged as bio-based alternatives for packaging and thermal insulation applications. India, the world’s largest producer of bananas, generates significant quantities of banana biomass (∼200 tons per hectare per year), much of which remains underutilized. Banana peduncle, the stalk that supports the fruit bunch, is one such underutilized biomass. In this study, banana peduncle fibers were used as the main substrate with *Pleurotus ostreatus* for the fabrication of MBCs. Banana peduncle fibers were mixed with wood shavings (10-50 wt%) to enhance the dimensional stability and structural integrity of the composites. The properties of developed MBCs such as density, shrinkage, moisture absorption, water absorption, morphology, compressive properties, and thermal conductivity were studied. The 90% banana peduncle fibers-10% wood shavings formulation showed the highest radial mycelial growth rate (7 mm/day). MBCs consisting of 100% banana peduncle fibers had volumetric shrinkage of 36%, while the incorporation of 30-50% wood shavings reduced shrinkage by approximately 17%. Among the formulations, MBCs containing 30% wood shavings had the highest compressive strength (4.82 MPa) and compressive modulus (1.78 MPa), whereas MBCs containing 50% wood shavings had the highest recovery (59.6%). In contrast, MBCs fabricated using 100% banana peduncle fibers had the lowest thermal conductivity (0.04 W/m. K). These results demonstrate that banana peduncle fibers are a promising lignocellulosic substrate for the development of MBCs for sustainable packaging and thermal insulation.

## 1. Introduction

Conventional materials such as expanded polystyrene (EPS) and polyurethane (PU) foams are extensively used for protective packaging and thermal insulation applications. However, these materials are non-biodegradable and pose significant environmental challenges after disposal. They also contribute to the generation of micro or nano-plastics, which is detrimental for humans, animals and the environment [1, 2]. According to the Central Pollution Control Board (CPCB), polystyrene accounts for ∼ 5% of the plastic waste generated across pilot Indian cities [3]. These environmental concerns have intensified the need for environment friendly, sustainable, and biodegradable alternatives [4].

In recent years, mycelium-based composites (MBCs) have emerged as a promising bio-based alternative for packaging and thermal insulation applications [4, 5]. MBCs are produced by cultivating fungal mycelium on lignocellulosic substrates. Their fabrication typically involves substrate preparation, fungal inoculation, and mycelial colonization, followed by heat treatment to terminate the fungal growth. During colonization, the fungal mycelium grows in and around the substrate particles, binding them together to form a solid composite [6, 7]. MBCs utilize renewable lignocellulosic residues, require less energy during fabrication and do not produce any toxic by-products, making them a sustainable alternative to conventional materials. The properties of the MBCs depend on the lignocellulosic substrate, fungal species and environmental conditions during fabrication [8].

Various lignocellulosic substrates such as wheat straw, rice straw, sawdust, woodchips, rice husk, hemp, flax, cotton, sugarcane bagasse, have been used for the fabrication of MBCs using fungal species such as *Ganoderma lucidum*, *Pleurotus ostreatus* and *Trametes versicolor*. Previous studies have demonstrated that both fungal species and type of substrate used influence the physical, mechanical and thermal properties of MBCs. For example, Elsacker et al. used hemp, flax, softwood, and straw as substrates together with *Trametes versicolor* for the fabrication of MBCs. Composites with chopped flax fibers had the highest compressive modulus (1.18 MPa), while composites with chopped hemp had a compressive modulus of 0.77 MPa. The thermal conductivity of composites with flax, hemp, and straw was in the range of 0.04-0.05 W/m. K [9]. Aiduang et al. utilized four fungal species (*Ganoderma fornicatum*, *Lentinus sajor-caju*, *Ganoderma williamsianum* and *Schizophyllum commune*) and three lignocellulosic substrates (rice straw, corn husk and sawdust) for the fabrication of MBCs. The MBCs had a water absorption of 105-208%. The MBCs had a density and compressive strength in the range of 190-340 kg/m^3^ and 0.25-1.87 MPa, respectively [10]. Peng et al. used sawdust, coir-pith, corn straw, rice straw and bagasse as substrates along with *Pleurotus ostreatus* for the fabrication of MBCs. Sawdust composites had the highest compressive strength (456.7 kPa), while corn straw composites had the lowest compressive strength (270 kPa). The moisture absorption at 90% RH was highest for corn straw composites (28%), lowest for coir-pith composites (22%) [11]. Wang et al. used 8 different fungal species and wheat straw as substrate for the fabrication of mycelium bio-foams. The densities of the composites ranged from 78 to 153 kg/m^3^. *Ganoderma lucidum* bio-foams had the highest shrinkage (29%) post-drying, while *Flammulina velutipes* bio-foams had the lowest shrinkage (2%). *Ganoderma lucidum* and *Pleurotus ostreatus* bio-foams showed compressive strength of 521 and 110 kPa at 65% strain and water absorption of 203% and 407%, respectively [12]. Zhong et al. used *Camellia oleifera* shell and *Ganoderma sessile* to produce MBCs, and the density and thermal conductivity of the developed composites were 295-297 kg/m^3^ and 0.041 W/m. K respectively [13]. These studies demonstrate that the selection of an appropriate lignocellulosic substrate and fungal species is critical for obtaining MBCs with desirable properties.

Banana peduncle is an underutilized agricultural residue in India. Banana is one of the most widely cultivated fruit crops in the world, with global production of approximately 139 million tons. India is the world’s largest banana producer, with an annual production of approximately 37.6 million tons (∼ 27% of global production) [14]. Thus, large quantities of banana biomass, including peduncles, pseudo stems, and leaves, are generated annually. Among the various banana biomass residues, the banana peduncle is the stalk that supports and connects the banana fruit bunch to the banana plant. Following the removal of banana fruits, the banana peduncle is typically discarded despite being rich in cellulose, hemicellulose, and lignin [15, 16]. In several studies, banana peduncle has been explored as a raw material to produce bio-adsorbents, biogas, paper and composites. For example, Muthusamy et al. utilized banana peduncle for the development of a bio-adsorbent for the removal of lead ions from industrial wastewater. The developed bio-adsorbent had a biosorption efficiency of 94% [17]. Fagbemigun et al. utilized banana peduncles to produce pulp and paper. The fabricated paper had a tensile index of 69.21 Nm/g [18]. Bishoyi et al. fabricated banana peduncle fiber-reinforced soy-resole composites. The tensile and flexural strengths of the composites were 43.5 MPa and 36.2 MPa, respectively [19]. However, to the best of our knowledge, banana peduncle has not been explored as a substrate for MBCs production.

Therefore, in this current study, MBCs were fabricated using banana peduncle fibers (BP) as the primary substrate and *Pleurotus ostreatus*. Wood shavings (WS) were added to BP in different ratios (10-50%) to enhance the dimensional stability and structural integrity of the composites. The radial and vertical growth of *Pleurotus ostreatus* mycelium on the substrate combinations were evaluated. Subsequently, the developed MBCS were characterized in terms of their density, shrinkage, moisture absorption, water absorption, morphology, compressive properties and thermal conductivity.

## 2. Materials and Methods

### 2.1 Materials

Fresh banana peduncles (Yelaki/Elaichi variety) were collected from Thiruvanmiyur market (12.9774° N, 80.2580° E), Chennai, India. The banana peduncles were cut into small pieces, dried in a hot air oven at 60°C for 2 days and ground in a mixer-grinder to produce the banana peduncle fibers. Wood shavings were collected from the Sree Mahalakshmi Timber shop, Velachery, Chennai, India and washed in hot water prior to use. The *Pleurotus ostreatus* mycelial culture was procured from ICAR-IIHR, India.

### 2.2 Characterization of the substrates (BP and WS)

BP and WS were viewed under a stereomicroscope (Nexius Zoom, Euromex, Microscopes Holland), and images of the fibers were captured with scale bar/magnifications. The lengths of BP/WS were measured using the Image Focus Alpha software, and average values (n=100) were reported. The carbon to nitrogen (C/N) ratio of the BP and WS was determined using an Elementar UNICUBE elemental analyzer, India. The TAAPI standard was used to calculate the water retention value of BP and WS substrates **(eq. (1))**. To make a homogeneous wet fiber pad, 5g of fibers were mechanically distributed in distilled water and filtered using a Buchner funnel. The wet fiber pad was then placed in a pre-weighed centrifuge tube with a porous filter to remove unbound water. The sample was centrifuged at 3000 g for 30 minutes. After centrifugation, the tube was removed, and the wet fibers were weighed on an electronic balance. The fibers in the tube were dried in a 105 ± 2°C oven until a steady weight was achieved. After drying, the sample was cooled in a desiccator and weighed again to determine oven dried fiber mass.

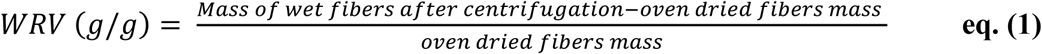

### 2.3 Growth of *Pleurotus ostreatus* on the four different BP-WS substrate combinations

The *Pleurotus ostreatus* culture was transferred onto sterile potato dextrose agar (PDA) petri-plates and incubated at 25°C for 2 weeks. After complete mycelial growth, the petri-plates were preserved at 4 °C until further use.

#### 2.3.1 Radial growth rate

To evaluate the radial growth, an agar plug (circular) containing the mycelial culture was placed at the center of a petri-dish containing a thin layer of substrate (BP100, BP90WS10, BP70WS30, BP50WS50). The petri-dishes were incubated at 25°C for 7 days. Photographs of the petri-dish were taken daily using a Samsung Smartphone, and the growth diameter (in mm) was measured using Image J. The measured growth diameter values were divided by two to obtain the growth radius values. For each of the substrate combinations, a graph of growth radius (mm) vs number of days was plotted **(Figure S3 (a))**. The slope of the linear region of the graph (day 2 to day 6) was used to calculate the radial growth rate (in mm/day) [12].

#### 2.3.2 Vertical growth rate

The colonization rate of *Pleurotus ostreatus* on BP and WS was estimated using the vertical growth rate. Test tubes were loosely filled with substrates (BP100, BP90WS10, BP70WS30, BP50WS50), and a circular inoculum was placed on the top of the substrate. The test tubes were incubated at 25 °C for 7 days, and the downward mycelial growth, referred to as vertical growth, was measured every day. Graphs were plotted between the vertical growth (in mm) vs. the number of days **(Figure S3 (b))**. The slope of the linear region of the graph was used to calculate the vertical growth rate (in mm/day) [20].

### 2.4 Development of the MBCs using BP-WS substrate combinations

The MBCs were fabricated using four different substrate combinations: BP100, BP90WS10, BP70WS30 and BP50WS50 (**Table 1**). The fabrication process involved four steps, namely substrate preparation, fungal inoculation, incubation and drying. The substrate combinations were soaked in water for 2 hours, after which the excess water was drained, and the moisture content was adjusted to 70%. The substrates were sterilized by autoclaving at 121 °C and 15 psi (Equitron). After cooling to room temperature, the sterilized substrates were inoculated with *Pleurotus ostreatus* spawn (20%; wet weight basis), thoroughly mixed and packed into cylindrical moulds. The molds were covered with cling film to prevent contamination and moisture loss during the incubation period. The inoculated substrates were incubated at 25°C for 2 weeks to allow mycelial colonization. After the incubation period, the colonized samples were removed from the moulds and dried at 70°C to remove the moisture, resulting in the formation of MBCs. The fabricated composites were designated as BP100, BP90WS10, BP70WS30 and BP50WS50 according to their respective substrate’s formulations.

**Table 1:** List of samples tested in this study.

| Sample code | Fungus used | Substrate used |
| --- | --- | --- |
| BP100 | *Pleurotus ostreatus* | Banana peduncle fibers (100%) |
| BP90WS10 | *Pleurotus ostreatus* | Banana peduncle fibers (90%) + Wood shavings (10%) |
| BP70WS30 | *Pleurotus ostreatus* | Banana peduncle fibers (70%) + Wood shavings (30%) |
| BP50WS50 | *Pleurotus ostreatus* | Banana peduncle fibers (50%) + Wood shavings (50%) |

### 2.5 Characterization of the banana peduncle fiber-mycelium-based composite

#### 2.5.1 Dry Density and Volumetric Shrinkage

The density and volumetric shrinkage percentage of the MBCs was calculated using the following formulas **(eq. (2) & eq. (3))**;

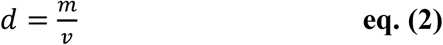

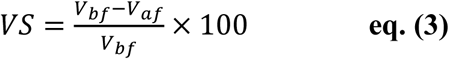

Where *d* is the density of MBC in g/cm^3^, *m* and *v are* the mass and volume of dried MBC, respectively; *VS* is the volumetric shrinkage percentage of MBC, and *V_bf_* is the volume of MBC before drying and *V_af_* is the volume of the MBC after drying.

#### 2.5.2 Moisture absorption

The MBCs were oven-dried at 105 °C, and their initial weight (*Wi*) was measured. The oven-dried MBCs were then placed in a humidity chamber (Climatic test chamber, GSC Global Make) at two different relative humidity (RH) conditions: 60% RH and 75% RH. The weight of the mycelium composite was measured every hour (*Wt*) until a constant weight was attained. The moisture absorption (%) was calculated using the equation below **(eq. (4))**. An analytical balance with an accuracy of 0.0001 g was used.

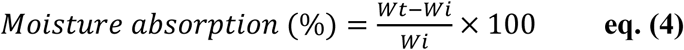

*Wi* is the initial dry weight of the composite, and *Wt* is the weight of the composite at any time t.

#### 2.5.3 Water absorption

The MBCs were oven-dried at 105°C, and their initial weights (*W_0_*) were measured. The MBCs were then immersed in containers filled with water. The weight of the MBCs was taken at 10,20,30,40,50,60,120,240,480,720 and 1440 minutes. At each time interval, the MBCs were removed from the container, and their post-immersion weight (*W_f_*) was measured. Water absorption percentage was calculated using the following formula **(eq. (5))**.

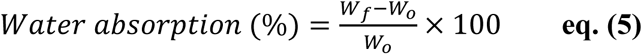

#### 2.5.4 Stereomicroscopy

The images of the top, bottom and cross-section of BP100, BP90WS10, BP70WS30 and BP50WS50 composites were taken using a stereomicroscope (Nexius Zoom, Euromex, Microscopes Holland) at different magnifications 10X, 20X and 35X.

#### 2.5.5 Scanning electron microscope

The substrates and the cross-section of the fabricated MBCs (BP100, BP90WS10, BP70WS30 and BP50WS50) were examined using a scanning electron microscope (Carl Zeiss Evo 18). Prior to imaging, all the samples were sputter-coated with a thin layer of gold (Quorum, SC7620, UK). Images of the samples were captured at different magnifications.

#### 2.5.6 Compression testing

Three cylindrical samples each of BP100, BP90WS10, BP70WS30 and BP50WS50 MBCs were tested using a universal testing machine (Dak System Inc Series 7200) equipped with a 5kN load cell. Tests were conducted at a displacement rate of 1 mm/min. The load-displacement curve was converted to a stress-strain curve using the following equations **(eq. (6) & eq. (7))**. The compressive strength was calculated at 70% strain. The compressive modulus was calculated from the slope of the linear region of the stress-strain curve (between 10-20% strain).

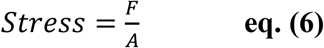

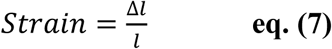

Where F is the compressive force in Newton, A is the cross-sectional area of the MBC in mm^2^, *l* is the original height of the MBC in mm and Δ*l* is the change in height in mm.

After compression to 70% strain, the compressive load was removed, and the samples were allowed to recover. The heights of the sample were measured at 1 min, 3 min and 30 min using a digital vernier caliper. The percentage recovery of height was calculated using the following formula.

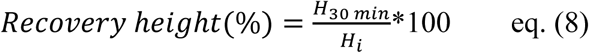

Where, *H*_30_ *_min_* is the recovery height at 30 min in mm and *H_i_* is the initial height before compression in mm.

#### 2.5.7 Thermal conductivity

The thermal conductivity of the BP100, BP90WS10, BP70WS30, and BP50WS50 was evaluated utilizing a Hot Disk Thermal Constants Analyzer (TPS 2500S, Hot Disk AB, Sweden) through the transient plane source (TPS) method using a Kapton 5465 F2 sensor.

## 3. Results and discussion

### 3.1 Properties of the lignocellulosic substrates

The average length of BP and WS used in this study were 8.72 ± 4.4 mm and 3.30 ± 1.6 mm, respectively **(Figure S1 (a) & (b)).** The C/N ratio is one of the important substrate characteristic and influences mycelial growth [21]. In this study, the C/N ratio of BP was 33.87, while that of WS was 682.14. The C/N ratios of the four substrate combinations used to fabricate MBCs were 34.07 (BP100), 38.7 (BP90WS10), 51.6 (BP70WS30) and 73.5 (BP50WS50). The C/N ratio of the substrate combinations increased with the increase in wood shaving percentage from 10 to 50%. The water retention capacity of lignocellulosic substrates is important for MBCs production, as it maintains the moisture required for mycelial growth and colonization. The water retention capacity of banana peduncle fibers was 1.518 g/g, while that of wood shavings was 0.909 g/g. BP contained 67.3 % holocellulose, 16.15% lignin and 7.30 % extractives **(Table 2)**. Holocellulose consists of both cellulose and hemicellulose fractions, which serve as the primary carbon source during fungal growth. The high holocellulose content along with moderate lignin content suggests that BP is a suitable substrate for *Pleurotus ostreatus* colonization. The radial and vertical growth of *Pleurotus ostreatus* on the substrate combinations is discussed in the subsequent sections.

**Table 2:** Composition of the substrates.

| Substrate | %<br>Carbon | %<br>Nitrogen | C/N ratio | Water<br>retention<br>value | %<br>extractives | %<br>lignin | %<br>holocellulose |
| --- | --- | --- | --- | --- | --- | --- | --- |
| BP | 38.16 | 1.12 | 33.87 | 1.518 g/g | 7.30 | 16.15 | 67.3 |
| WS | 49.28 | 0.07 | 682.14 | 0.909 g/g | - | - | - |

### 3.2 Radial and vertical growth of *Pleurotus ostreatus* mycelium on different substrate combinations

Photographs were captured on days 0, 2, 4 and 6 to evaluate the radial growth of *Pleurotus ostreatus* mycelium on different substrate combinations **(Figure 1 (a-d))**. On day 0, only the *Pleurotus ostreatus* inoculum (agar plug) was present in all the samples. By day 2, *Pleurotus ostreatus* mycelium started to grow radially from the edge of the inoculum. By day 4, the mycelium continued to expand radially across the substrates. After 6 days of growth, the mycelium completely covered the substrate surface in the case of BP100 and BP90WS10. In contrast, BP70WS30 and BP50WS50 exhibited comparatively lower surface mycelial coverage. The radial growth rate of *Pleurotus ostreatus* on BP100, BP90WS10, BP70WS30 and BP50WS50 was 6.82 ± 0.15, 7 ± 0.15, 6.57 ± 0.08, and 4.92 ± 0.08 mm/day, respectively **(Figure 1(e))**. The substrate composition influences the mycelial growth rate, because the fungal hyphae are in direct contact with the substrate and obtain the nutrients required for growth from it [22]. The highest radial growth rate was observed for BP90WS10, while BP50WS50 had the lowest radial growth rate. The lowest radial growth rate in BP50WS50 may be due to its high C/N ratio (73.5), which may have resulted in nitrogen limitation [23]. Schoder et al. reported a radial expansion rate of 4.33 mm/day for *Pleurotus ostreatus* grown on cotton fibers [22]. Sangkawanna et al. reported the radial growth rate of *Pleurotus pulmonarius* on water hyacinth/sawdust substrates to be 7.2-10.8 mm/day [24].

**Figure 1:**
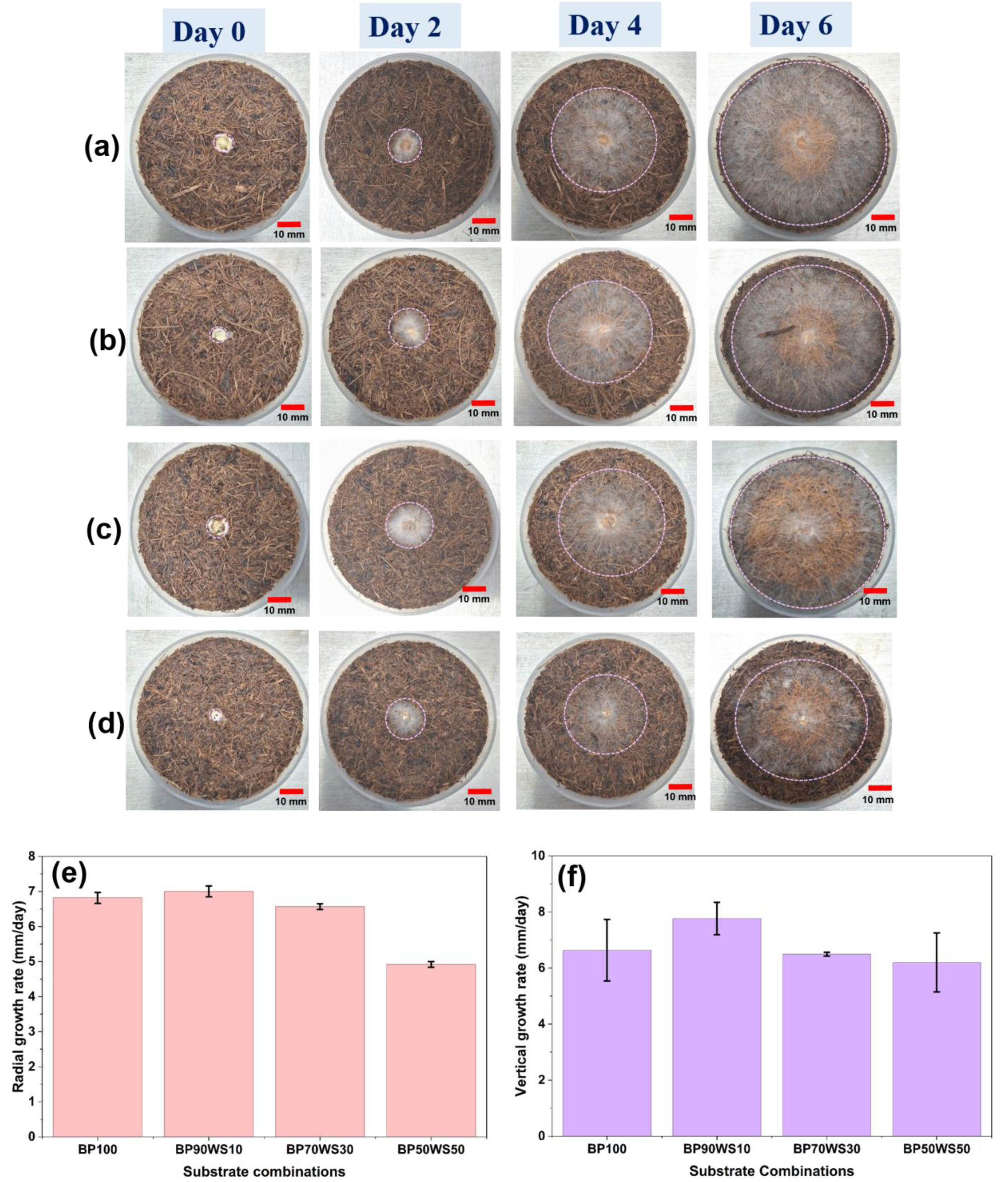
Photographs of the Radial growth of *Pleurotus ostreatus* mycelium on different substrate combinations (a) BP100, (b) BP90WS10, (c) BP70WS30 and (d) BP50WS50. Graphs showing the (e) radial growth rate and (f) vertical growth rate of the *Pleurotus ostreatus* mycelium on different substrate combinations

The vertical growth rate of *Pleurotus ostreatus* on the substrate combinations BP100, BP90WS10, BP70WS30, and BP50WS50 was 6.63 ± 1.1, 7.77 ± 0.58, 6.49 ± 0.07 and 6.2 ± 1.05 mm/day, respectively **(Figure 1(f))**. Among the substrate combinations, BP90WS10 had the highest vertical growth rate, while BP50WS50 had the lowest vertical growth rate. This shows that the BP90WS10 substrate promoted faster mycelial penetration and colonization. The photographs showing the downward growth of mycelium on different substrate combinations are shown in **Figure S2 (a-d)**. Zervakis et al. reported the colonization rate of *Pleurotus ostreatus* on wheat straw, peanut shells, poplar sawdust, oak sawdust, corn cobs and olive press cake to be 6.2, 8.6, 7.5, 6, 7.1 and 3.4 mm/day, respectively [20].

### 3.3 Physical Properties of MBCs

In this study, the density of the fabricated MBCs was in the range of 0.283-0.304 g/cm^3^ **(Figure 2 (a))**. The densities of BP100, BP90WS10, BP70WS30 and BP50WS50 MBC were 0.30 ± 0.03 g/cm^3^, 0.29 ± 0.02 g/cm^3^, 0.28 ± 0.02 g/cm^3^ and 0.30 ± 0.01 g/cm^3^ respectively. Appels et al. reported the densities ranging from 0.13 - 0.39 g/cm^3^ [25], while Peng et al. and Cai et al. reported densities ranging from 0.24 - 0.33 g/cm^3^ [11] and 0.25 - 0.37 g/cm^3^ [26] respectively. The density values obtained in this current study were comparable to those reported in the literature.

**Figure 2:**
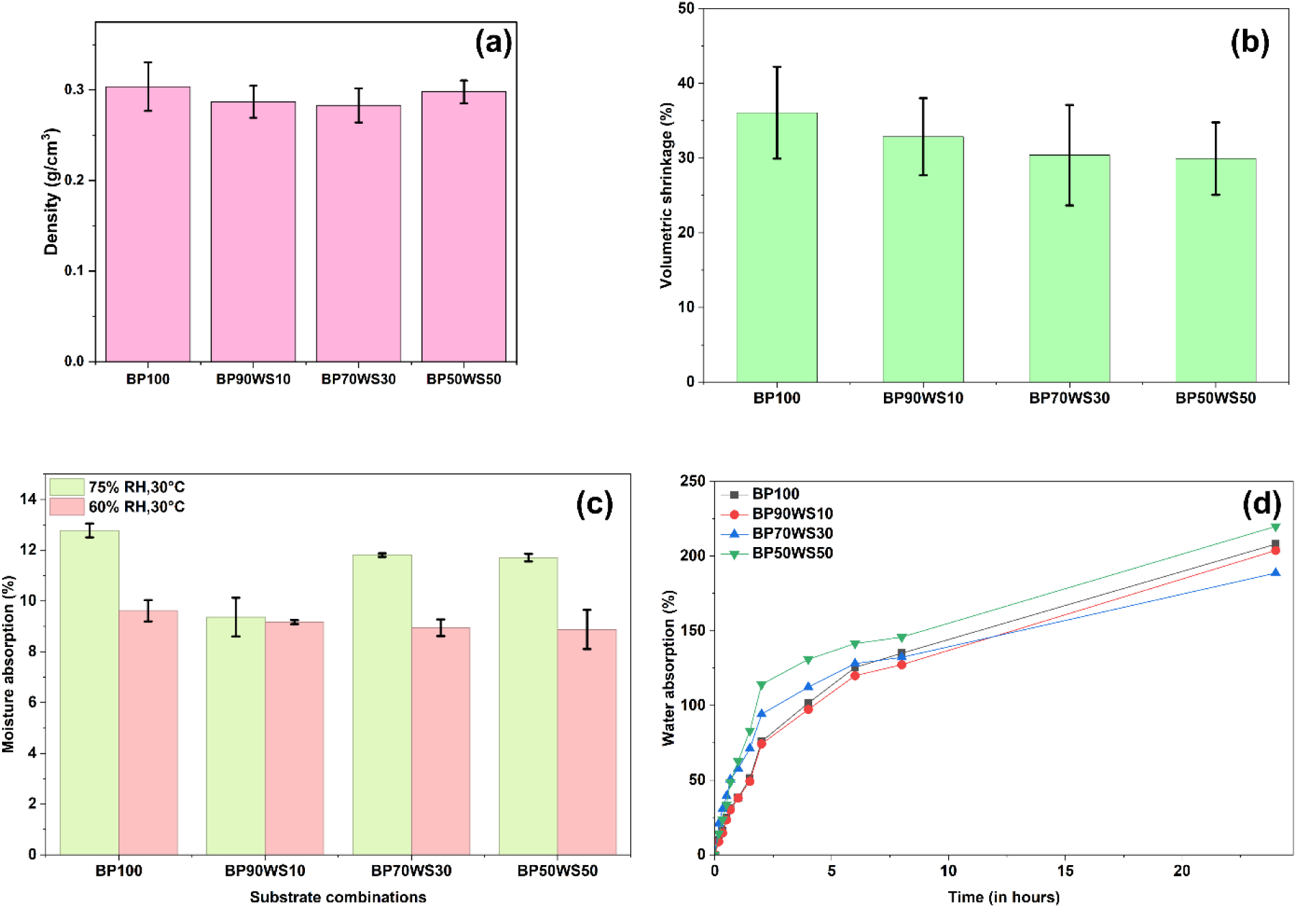
Represents the (a) density, (b) volumetric shrinkage (%) during drying (c) moisture absorption (%) at 60% and 75% RH and (d) water absorption of the MBCs.

The volumetric shrinkage of the fabricated MBCs ranged from 29.9% - 36.1% **(Figure 2 (b))**.The volumetric shrinkage of BP100, BP90WS10, BP70WS30, and BP50WS50 were found to be 36.1 ± 6.2 %, 32.9 ± 5.1 %, 30.4 ± 6.7 %, and 29.9 ± 4.8 % respectively. BP100 exhibited the highest shrinkage, while BP50WS50 and BP70WS30 exhibited the lowest shrinkage. Volumetric shrinkage decreased with an increase in the wood shaving percentage in the composites. This decrease might be due to the lower water retention capacity of WS (0.909 g/g) compared to BP (1.518 g/g), which could influence the dimensional stability of the composites. Wang et al. reported average shrinkage of 2 - 29% [12], while Elsacker et al. and Aiduang et al. reported average shrinkage of 9 - 20% [9] and 8 - 16% [10], respectively. The percentage shrinkage of the MBCs in this current study was higher than that reported in the literature. The higher volumetric shrinkage observed in this present study, may be due to the differences in the lignocellulosic substrate used compared with previous studies.

The equilibrium moisture absorption of the fabricated MBCs, measured after 8 hours of exposure, ranged from 8.9 - 9.6 % at 60% RH, and 9.4 - 12.8 % at 75% RH **(Figure 2 (c))**. The moisture absorption of the MBCs increased with an increase in relative humidity. At 60% RH, the moisture absorption of BP100, BP90WS10, BP70WS30, and BP50WS50 was 9.6 ± 0.4 %, 9.2 ± 0.1 %, 8.9 ± 0.3 % and 8.9 ± 0.8 %, respectively. At 60 % RH, the moisture absorption of the four different MBCs were similar to each other. At 75% RH, the moisture absorption of BP100, BP90WS10, BP70WS30, and BP50WS50 was 12.8 ± 0.3 %, 9.4 ± 0.7 %, 11.8 ± 0.1 % and 11.7 ± 0.2 %, respectively. At 75% RH, BP90WS10 had the lowest moisture absorption among the other composites. The graphs between moisture absorption percentage and time (hours) are depicted in **Figure S4 (a-b)**. The moisture absorption of the MBC increased rapidly within the first 2 - 4 hours, followed by a slower increase until equilibrium was reached after ∼ 8 hours. These values are comparable to those reported in previous studies. Qiu et al. reported the moisture absorption values of 4.49 - 7.21% at 60% RH and 10.68 - 15.42% at 80% RH, 40°C [27]. Appels et al. reported values of 3.15 - 8.22% at 60% RH and 7.57 - 11.63% at 80% RH, 40°C [25].

The water absorption of the MBCs after 24 hours of immersion in water ranged from 188-219% **(Figure 2(d))**. BP100, BP90WS10, BP70WS30 and BP50WS50 had water absorption values of 208.1 ± 18.3 %, 203.8 ± 40.7 %,188.7 ± 40.8 % and 220 ± 6.7 % respectively. Despite differences in substrate compositions, the 24 hour water absorption was comparable among all the four composites. During the initial 120 minutes of immersion, the water absorption rate differed among the four substrate combinations. BP50WS50 had the highest initial water absorption rate (0.91%/min), while BP90WS10 had the lowest (0.57%/min). While BP100 and BP70WS30 had an initial water absorption rate of 0.58%/min and 0.69%/min, respectively. The higher initial water absorption rates of BP50WS50 and BP70WS30 may be due to the reduced surface mycelial colonization and a greater exposure of fibres on the outer surface of the composites. In contrast, BP100 and BP90WS10 had a dense white mycelial outer layer, which may have delayed the intial penetration of water (**Figure 3**). Cai et al. reported water absorption values ranging from 190-396% for MBCs fabricated using agricultural straws and *Ganoderma lucidum* mycelium [26]. Similarly, Wang et al. reported water absorption of 262-407 % for MBCs made up of wheat straw and *Pleurotus ostreatus* mycelium [12]. The water absorption values obtained in this study are comparable to that of literature.

**Figure 3:**
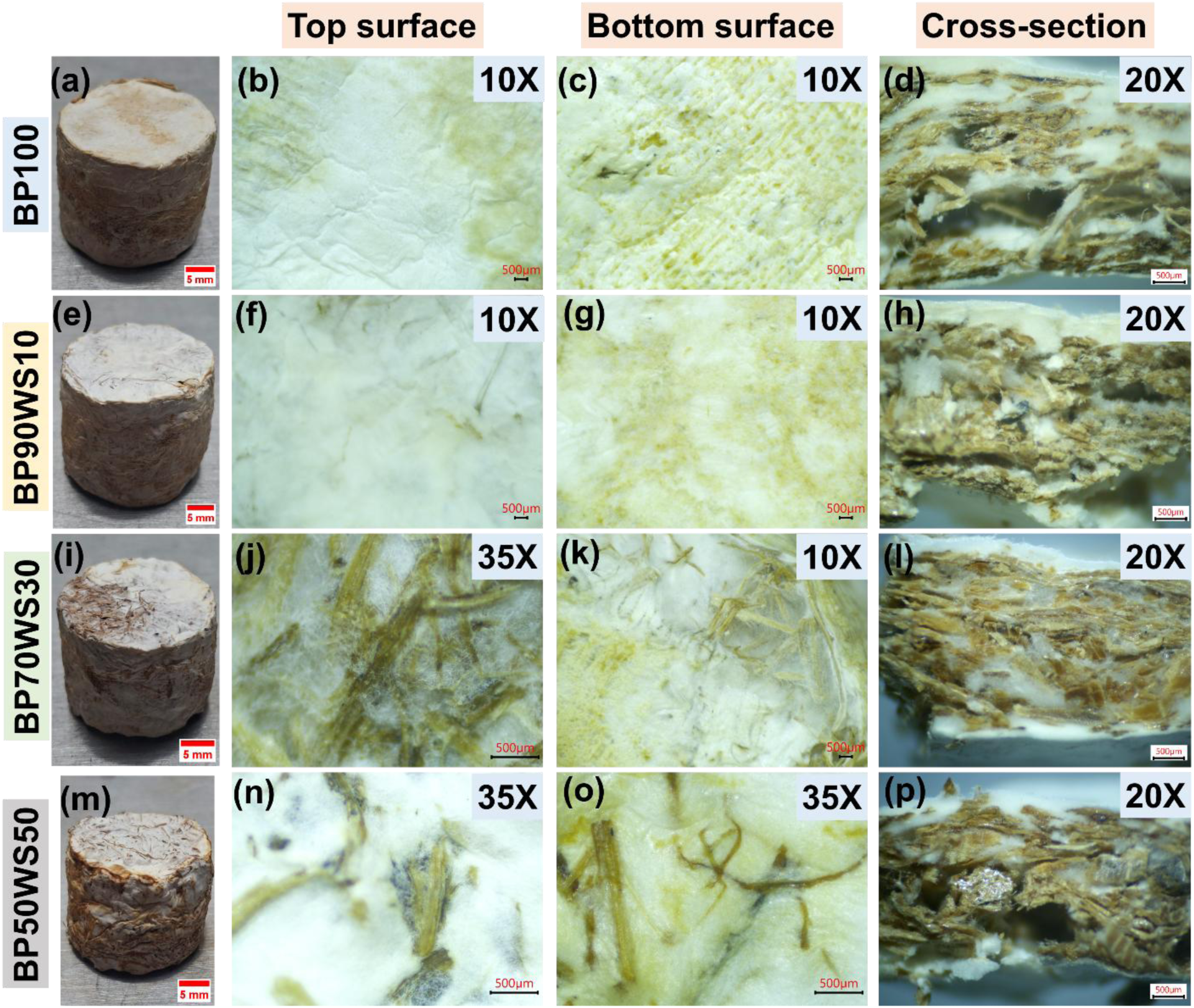
Representative photographs and stereomicroscopic images of the top surface, bottom surface and cross-section of MBCs (a-d) BP100, (e-h) BP90WS10, (i-l) BP70WS30 and (m-p) BP50WS50. The top and bottom surfaces were imaged at10X or 35 X, while cross-sections were imaged at 20 X.

### 3.4 Morphological characterization of MBCs

#### 3.4.1 Stereomicroscopy

The photographs of the cylindrical MBCs fabricated in this study are shown in **Figure 3(a, e, i, m)**. The stereomicroscopic images were taken to understand the external surface morphology of the MBCs. **Figure 3** shows the stereomicroscopic images of the top layer, bottom layer and cross-section of the MBCs. In the case of BP100 **(Figure 3 (b) & (c))** and BP90WS10 **(Figure 3 (f) & (g))**, the *Pleurotus ostreatus* mycelium completely covered the substrate, forming a dense white mycelial layer. The formation of this dense white top layer indicated uniform surface mycelial colonization. In the case of BP70WS30 **(Figure 3 (j) & (k))** and BP50WS50 **(Figure 3 (n) & (o))**, the substrate fibers were visible on the top and bottom layers, indicating reduced surface mycelial colonization. The reduced surface colonization observed for BP70WS30 and BP50WS50 may be due to their higher wood shaving content (30-50%) and higher C/N ratios of the substrate. The C/N ratio of BP50WS50 (73.5) exceeded the reported optimum range (45-60) for *Pleurotus ostreatus* growth, which may have contributed to the reduced surface mycelial colonization [21].

#### 3.4.2 Scanning electron microscopy (SEM)

##### 3.4.2.1 SEM images of lignocellulosic substrates

The banana peduncle fibers had a rough and irregular surface, with the presence of surface grooves **(Figure 4 (a))** [28]. The cross-section of the banana peduncle fiber bundle exhibited a porous structure with the presence of multiple hollow cavities known as lumens **(Figure 4 (b))**. Similar porous structures have been reported by Soraisham et al. [29], Anafack et al. [30], and Ruangnarong et al. [31]. The presence of these lumen structures might enhance the moisture absorption and thermal insulation properties of the banana peduncle fibres [29, 30, 32]. The major and minor lumen diameters were 24.17±4.36 µm and 13.11±3.92 µm, respectively (calculated via Image J). Ferdous et al. reported the lumen diameter of banana peduncle fibers to be 21.7 µm [33]. The wood shavings had a rough surface morphology **(Figure 4(c))**, with the presence of small porous structures (pores) **(Figure 4(d)).** The pore diameter was 1.77 ± 0.505 µm (calculated via Image J). The presence of similar pores in wood shavings has also been reported by Benchouaf et al. [34].

**Figure 4:**
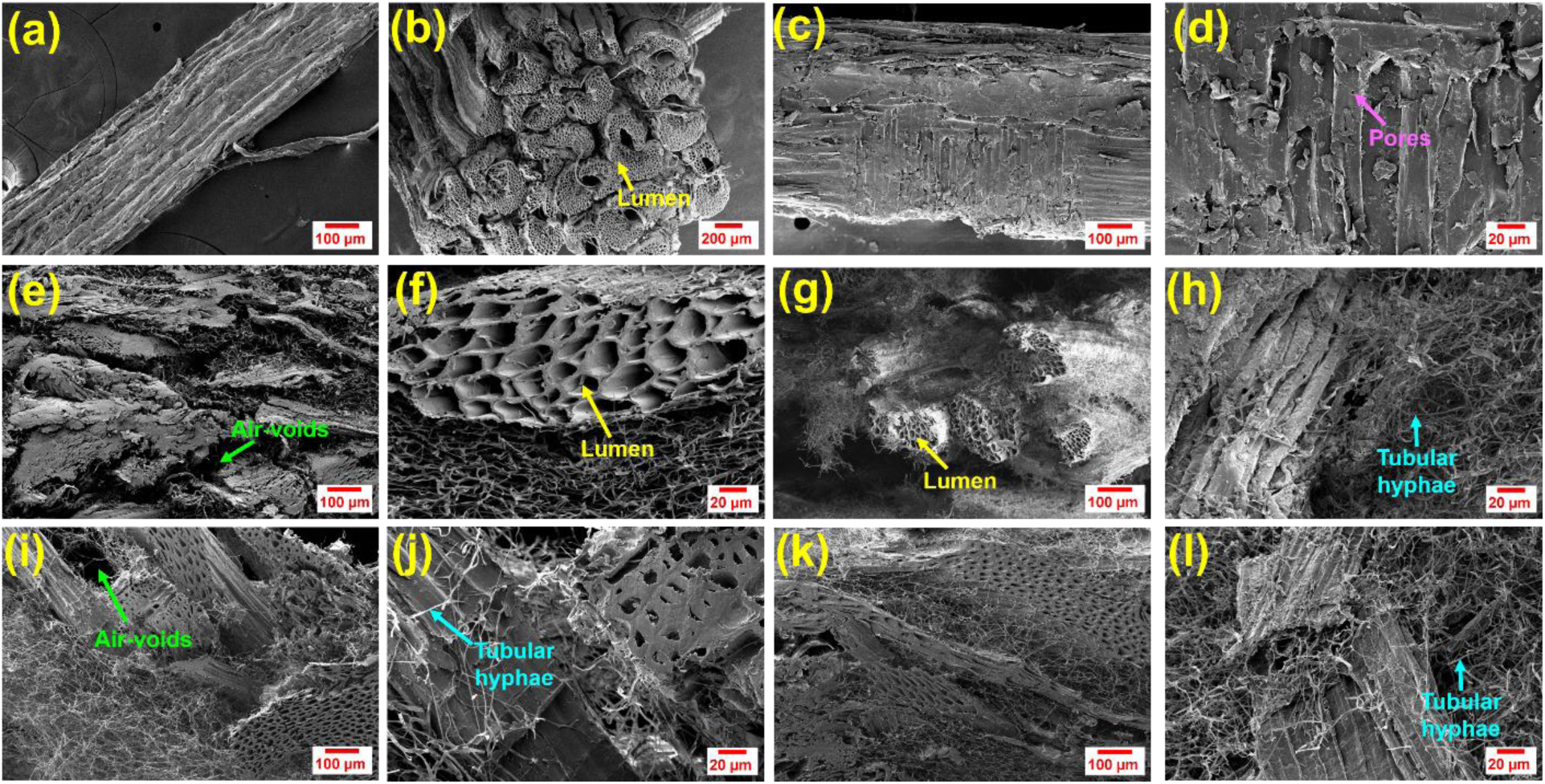
SEM images of the lignocellulosic substrates and MBCs. (a) Surface morphology of banana peduncle fibers, (b) cross-section of banana peduncle fiber bundle showing lumen structures, (c-d) surface morphology of wood shavings. SEM images of the cross-section of (e-f) BP100, (g-h) BP90WS10, (i-j) BP70WS30, (k-l) BP50WS50 MBCs at magnifications of 250 X and 1.00 KX.

##### 3.4.2.2 SEM images of cross-section of MBCs

The SEM images of the composite’s internal structure are shown in **Figure 4 (e-l)**. The SEM images indicate successful mycelium growth inside the composites. The SEM images showed the presence of lignocellulosic fibers, tubular hyphae, air voids and lumen structures (hollow cavities). The fungal hyphae were observed growing on and around the fiber surfaces **(Figure 4 (g), (h), (j), (l))**. The cross-section of BP100 and BP90WS10 exhibited a relatively homogenous internal morphology dominated by banana peduncle fibers, whereas BP70WS30 and BP50WS50 exhibited a more heterogeneous morphology due to increased presence of wood shavings. Similar findings were also reported by Aiduang et al. Appels et al. and Wang et al. [10, 12, 25].

### 3.5 Mechanical properties of MBCs

In this study, the compressive strength, compressive modulus and recovery of the MBCs were evaluated. The stress-strain curves of the MBCs are shown in **Figure 5 (a)**. All four composites (BP100, BP90WS10, BP70WS30 & BP50WS50) exhibited similar stress-strain curves consisting of an initial linear elastic region followed by a densification region. In the linear elastic region, the stress increased linearly with strain. At higher strains, stress increased rapidly due to the compaction of the composite structure and increased contact between the substrate particles. The average compressive strength at ∼ 70 % strain ranged from 4.07 to 4.82 MPa. Among the formulations, BP70WS30 had the highest average compressive strength (4.82 ± 0.69 MPa), followed by BP100 (4.77 ±1.67 MPa), BP50WS50 (4.27 ± 1.41 MPa), and BP90WS10 (4.07 ± 1.57 MPa). The average compressive strengths at 70% strain were almost similar because the composites had entered the densification region. Previous studies have reported that the compressive strength of MBCs is influenced by the type of lignocellulosic substrate used during their fabrication [11].

**Figure 5:**
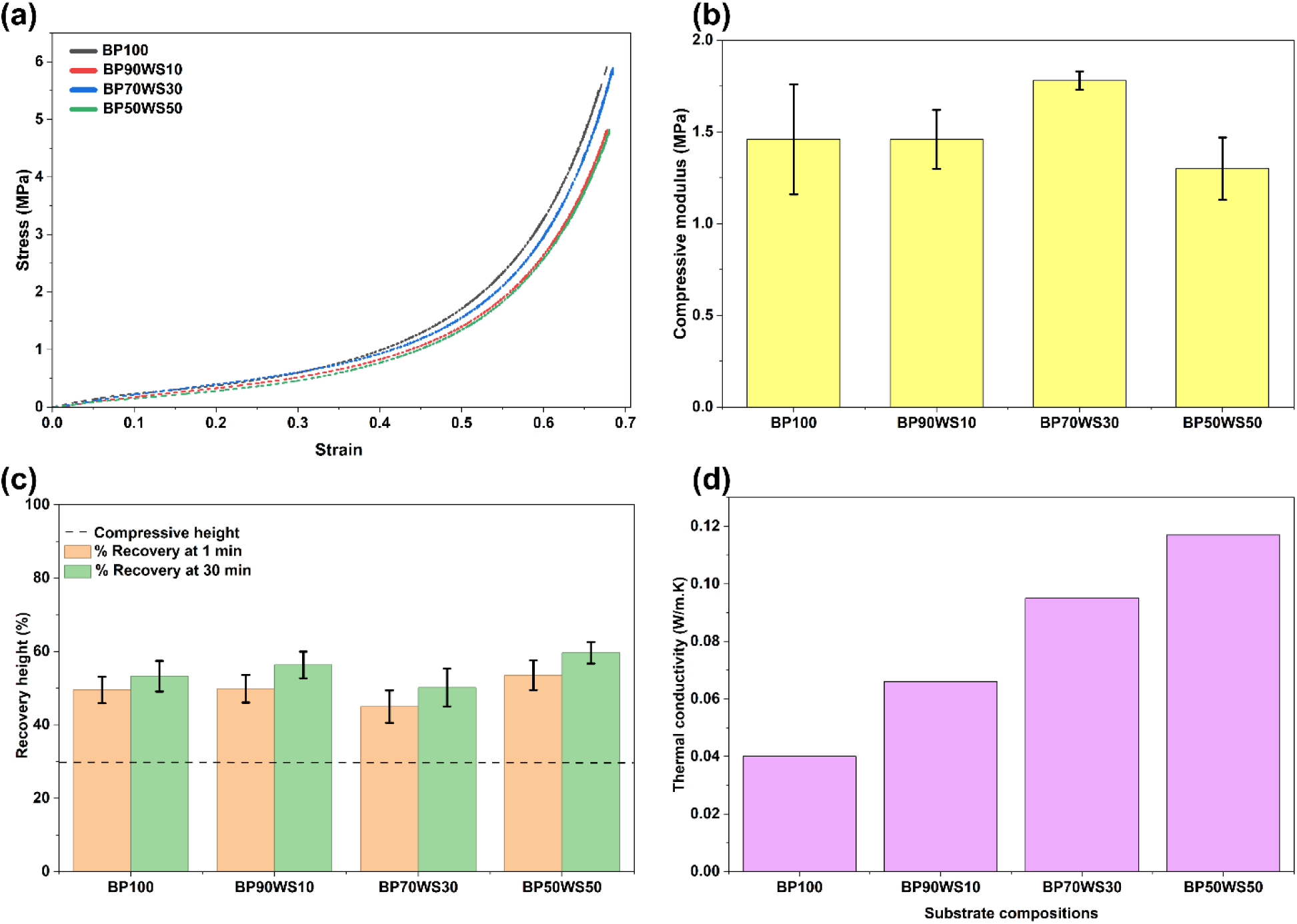
Represents (a) Stress-strain curves (b) compressive modulus, (c) % recovery after compression test and (d) thermal conductivity of MBCs

The compressive modulus (calculated from the slope of the stress-strain curve between 10-20% strain) was in the range of 1.30 to 1.78 MPa **(Figure 5 (b))**. BP70WS30 had the highest compressive modulus (1.78 ± 0.05 MPa), while BP50WS50 had the lowest compressive modulus (1.30 ± 0.17 MPa). Both BP100 and BP90WS10 exhibited similar compressive modulus of 1.46±0.3 MPa and 1.46± 0.16 MPa, respectively. BP70WS30 was the stiffest among all the composites. Elsacker et al. reported the compressive Young’s modulus of 1.18 MPa for MBCs fabricated using chopped flax fibers and *Trametes versicolor* [9].

As shown in **Figure 4 (e-l)**, the BP and WS substrate particles were interconnected via a dense mycelial network. This interconnected mycelial network likely contributed to the partial recovery of the composites after compression testing. Upon unloading, the elastic deformation recovered immediately, whereas a fraction of plastic deformation remained. However, complete recovery was not achieved, mostly due to irreversible deformation of the mycelial network, fiber rearrangement and collapse of air-voids during compression [35]. The recovery (%) of the MBCs after the compression tests is shown in **Figure 5 (c)**. The composites recovered 45-53% of their original height within 1 minute. The recovery increased slightly to 48 - 55 % after 3 minutes and 50-60% after 30 minutes. These results indicate that most of the height recovery occurred within the first minute after unloading. Among the four composites, BP50WS50 had the highest recovery (59.66 ± 2.95%), while BP70WS30 had the lowest recovery (50.17 ± 5.17 %). BP100 and BP90WS10 had intermediate recovery of 53.26 ± 4.19 and 56.35 ± 3.62 %, respectively. The comparatively lower recovery of BP70WS30 may be due to its higher compressive modulus. Wang et al. reported a recovery of 73-80% for *Pleurotus ostreatus* bio-foams and 51-55% for *Ganoderma lucidum* bio-foams [12].

### 3.6 Thermal conductivity of MBCs

In general, materials with thermal conductivity less than 0.07 W/m. K are generally considered to be good insulators [36]. Commonly used thermal insulation materials such as EPS and glass wool have a thermal conductivity of 0.029-0.040 W/m. K [37]. In this study, the thermal conductivity of the MBCs ranged from 0.040 to 0.117 W/m. K **(Figure 5 (d))**. BP100 had the lowest thermal conductivity (0.040 W/m. K), while BP50WS50 had the highest thermal conductivity (0.117 W/m. K). The thermal conductivity of BP90WS10 and BP70WS30 was 0.066 W/m. K and 0.095 W/m. K, respectively. Thermal conductivity increased with increasing wood shavings content in the composites. The thermal conductivity of MBCs depend on its substrate composition and microstructure [37, 38]. As observed in SEM images **(Figure 4 (b))**, banana peduncle fibers contain porous lumen structures which can entrap air and reduce heat transfer through the composites. Since, BP100 and BP90WS10 contained a higher percentage of banana peduncle fibers, they exhibited lower thermal conductivity. Similar observations have been reported in literature. Zhang et al. reported that mycelium composites made using birch sawdust had a higher thermal conductivity (0.05-0.074 W/m. K) than those fabricated using poplar sawdust (0.04-0.05 w/m. K) [36]. Zhang et al. in their study, produced a mycelium composite insulation brick using rye berries and *Pleurotus ostreatus*. The density and thermal conductivity of the mycelium insulation brick were 599 kg/m^3^ and 0.069 W/m. K, respectively [39]. Bonga et al. used coffee silver skin flakes and *Pleurotus ostreatus* to fabricate mycelium composites for thermal and acoustic insulation. The thermal conductivity of the mycelium composite was 0.03-0.04 W/m. K [40].

## 4. Conclusion

MBCs were successfully fabricated using banana peduncle fibers as the primary substrate and *Pleurotus ostreatus* mycelium. Wood shavings were added to banana peduncle fibers at different proportions (10-50%) to enhance the dimensional stability and structural integrity of the composites. The radial (7 mm/day) and vertical growth (7.77 mm/day) of *Pleurotus ostreatus* mycelium was highest on substrates containing 10% wood shavings. The fabricated MBCs exhibited densities of 0.28 - 0.30 g/cm^3^, volumetric shrinkage of 29.9 - 36 %, compressive strength of 4.07 - 4.82 MPa, compressive modulus of 1.3 - 1.78 MPa and thermal conductivity of 0.04 - 0.117 W/m. K. Overall, the results demonstrate that banana peduncle fibers are a promising lignocellulosic substrate for the fabrication of MBCs. The substrate combinations consisting of 90% banana peduncle fibers and 10% wood shavings provided the most balanced combination of rapid mycelial growth, good mechanical performance (compressive strength of 4.07 MPa, compressive modulus of 1.46 MPa and height recovery after compression of 56.35%) and low thermal conductivity (0.066 W/m. K), making it a promising candidate for sustainable packaging and thermal insulation applications.

## 5. Acknowledgments

The experimental work is funded through the NFIG grant of IIT Madras and Ministry of Education, Goverment of India.

