## supplementary data for "Upcycling banana peduncle fibers into mycelium-based composites for sustainable packaging and thermal insulation"

### Supplementary file

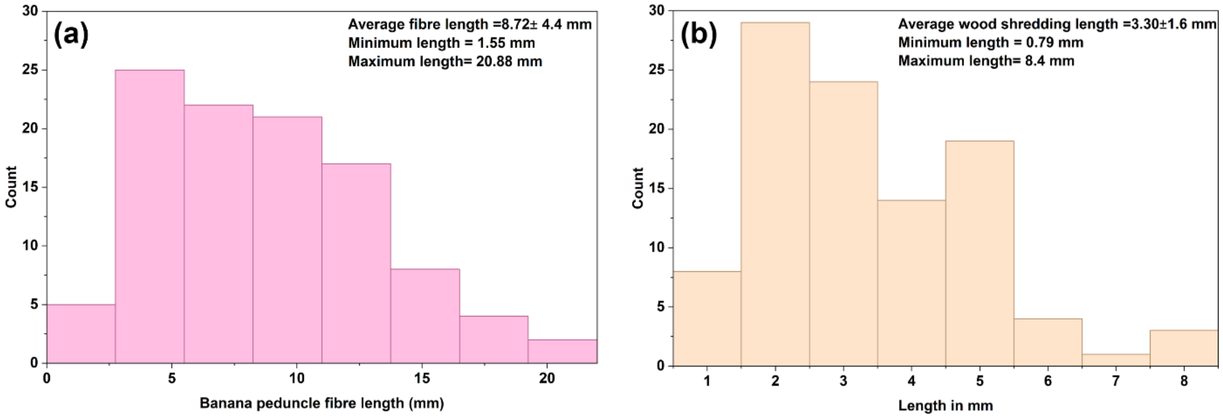


**Figure S1**: Average length distribution of (a) Banana peduncle fibers (BP) and (b) Wood shavings (WS)

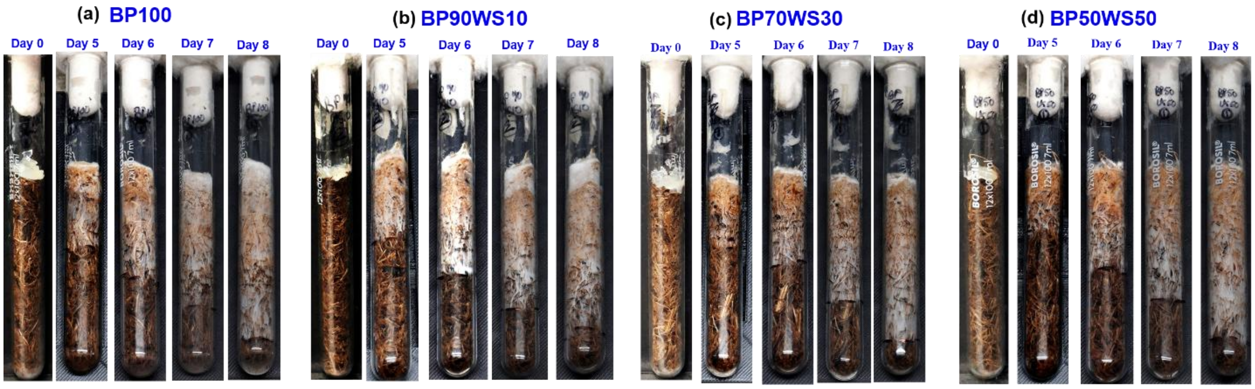


**Figure S2**: Photographs of the vertical growth of Pleurotus ostreatus mycelium on different substrate combinations (a) BP100, (b) BP90WS10, (c) BP70WS30 and (d) BP50WS50

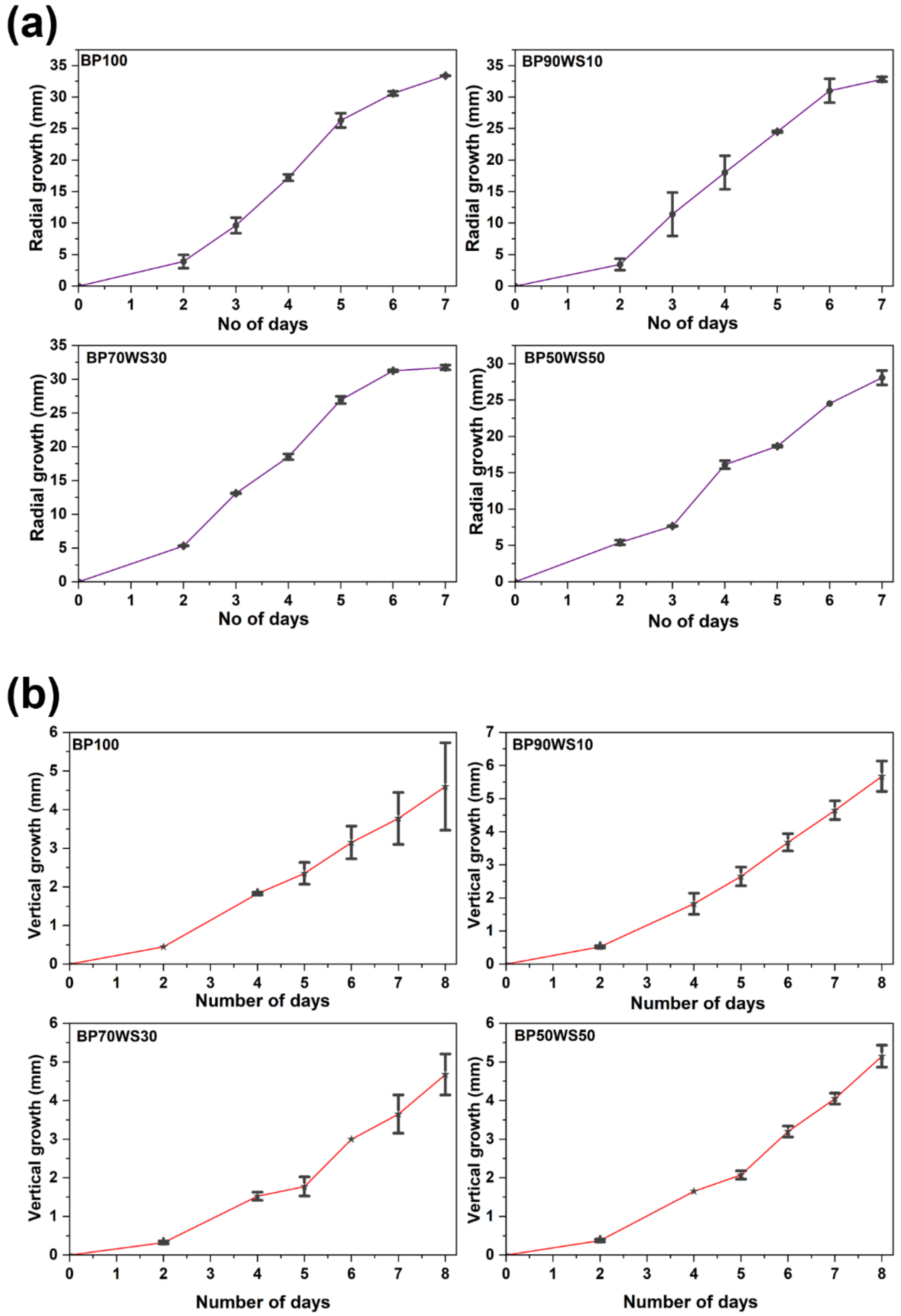


**<u>Figure S3</u>:** Graphs representing the (a) radial and (b) vertical growth of *Pleurotus ostreatus* on the four different substrate combinations

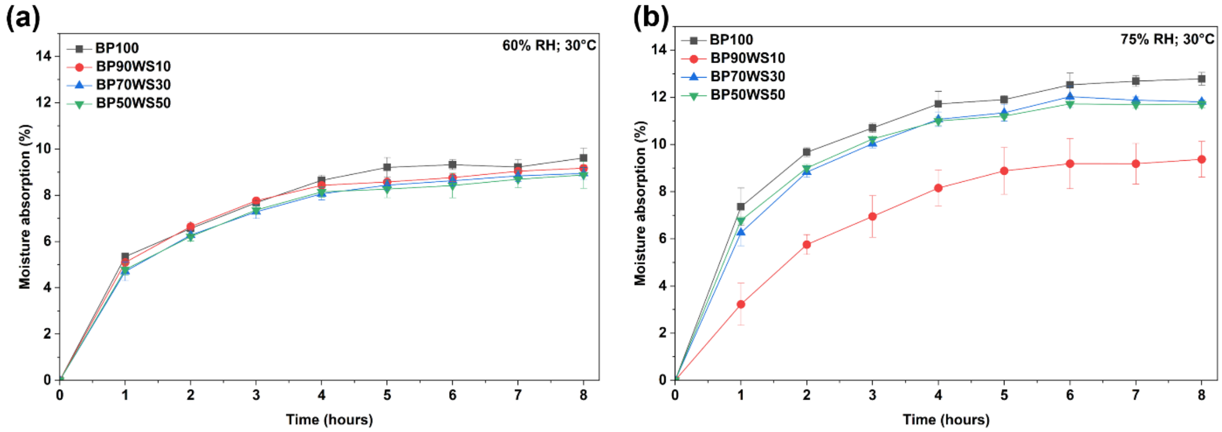


**<u>Figure S4</u>:** Graphs representing the moisture absorption of mycelium-based composites with time at (a) 60% RH and (b) 75 % RH.
